# Non-invasive MRI of Blood-Cerebrospinal Fluid-Barrier Function: a Functional Biomarker of Early Alzheimer’s Disease Pathology

**DOI:** 10.1101/2024.03.06.583668

**Authors:** Charith Perera, Renata Cruz, Noam Shemesh, Tania Carvalho, David L. Thomas, Jack Wells, Andrada Ianus

**Affiliations:** UCL Centre for Advanced Biomedical Imaging, Division of Medicine, University College London, London, United Kingdom; Champalimaud Research, Champalimaud Foundation, Lisbon, Portugal; Neuroradiological Academic Unit, Department of Brain Repair and Rehabilitation, UCL Queen Square Institute of Neurology, London, United Kingdom; Dementia Research Centre, UCL Queen Square Institute of Neurology, London, United Kingdom; Wellcome Centre for Human Neuroimaging, UCL Queen Square Institute of Neurology, University College London, London, United Kingdom

**Keywords:** Alzheimer’s disease, choroid plexus, blood CSF barrier, brain perfusion, arterial spin labelling MRI, 3xTg mouse model

## Abstract

**INTRODUCTION:** Choroid plexus (CP) dysfunction is thought to contribute to toxic protein build-up in neurodegenerative disorders, including Alzheimer’s disease (AD). However, the dynamics of this process remain unknown, mainly due to the paucity of in-vivo methods capable of assessing CP function.

**METHODS:** Here, we harness recent developments in Arterial Spin Labelling MRI to measure water delivery across the blood cerebrospinal fluid barrier (BCSFB) as a proxy for CP function, as well as cerebral blood flow (CBF), at different stages of AD progression in the widely used triple transgenic mouse model (3Tg), which recapitulates aspects of disease pathology.

**RESULTS:** Total BCSFB-mediated water delivery is significantly higher in 3Tg mice (>50%) from 8 weeks (preclinical stage), while tissue parameters such as CBF and T1 are not different between groups at all ages.

**DISCUSSION:** Our work shows changes in BCSFB function in the early stages of AD, providing a novel biomarker of pathology.

## 1. Introduction

The choroid plexus (CP) or blood-cerebrospinal fluid-barrier (BCSBF) is a unique functional tissue that pervades the brain’s fluid-filled ventricles. It is composed of a tight epithelium and stroma fed by a dense network of permeable capillaries, giving rise to a rate of arterial perfusion 5-10× greater than gray matter [1]. As the primary source of cerebrospinal fluid (CSF) replenishment in the central nervous system and a key site of clearance to the circulation [2], healthy BCSFB function is likely to be deeply entwined with the workings of CSF-mediated brain clearance processes such as the glymphatic system [3].

Dysfunction of the CP-CSF-axis is thought to contribute to the build-up of toxic proteins that are one of the hallmarks of Alzheimer’s disease (AD) [4–7], with CSF biomarkers showing promise in the diagnostic workup of AD [8]. Moreover, post-mortem studies [9,10] and preclinical ex-vivo studies [11] have shown AD-related differences in CP gene expression and morphology. To date, however, our understanding of CP function and its dynamics over the course of the disease has been restricted by a lack of non-invasive measurement techniques meaning that, despite its unique physiological significance, there is relatively little *in-vivo* data describing CP function in the healthy and diseased brain.

Recently, several imaging studies have taken an interest in mapping CP changes related to AD pathology, studying alterations in various aspects of morphology and function. For instance, in patients with AD, structural MRI data has shown an increase in CP volume with disease severity [12–14], arterial spin labelling (ASL) MRI and dynamic contrast enhanced (DCE) MRI have shown alterations in CP blood flow [14] and permeability to contrast agent, respectively [12], while dynamic PET has revealed a reduction in CSF clearance of tau and amyloid tracers [15–17]. Many of these techniques make use of exogenous tracers that might not necessarily reflect the BCSFB-mediated transport [18] and can have limitations for clinical usage and longitudinal studies. The general principle of ASL is to label a bolus of intravascular blood water using RF pulses (ie. without exogenous tracers), and subsequently measure the difference between control and labelled images at different inflow times, thereby permitting the non-invasive quantification of rates of water delivery to the tissue as an estimate of tissue perfusion.

To measure CP function in-vivo without using external tracers, a novel MRI approach based on Arterial Spin Labelling (ASL) at ultra long echo time (TE) was recently proposed [19] to measure the rate of delivery of labelled arterial blood water into ventricular CSF across the BCSFB. To date, this approach has been employed to study changes in BCSFB function in rodent models of ageing [19], pharmacomodulation [20] and systemic hypertension [21]. Recent work has also demonstrated the feasibility of this translational approach to capture the dynamic exchange of labelled blood-water into the CSF in the human brain [22].

Here, we harness ASL MRI to investigate, in-vivo, the alterations of CP function due to AD pathology, its dynamics with disease progression as well as compare it to cerebral perfusion. To this end, we capture MRI measurements of cerebral blood flow and BCSFB water delivery at different disease stages, together with behavioural testing and histopathology analysis in the widely employed triple transgenic mouse model of AD (3Tg-AD), which recapitulates both amyloid and tau pathology. To our knowledge this represents the first investigation of choroid plexus function in AD using non-invasive, translational imaging techniques, which may provide a novel and sensitive biomarker of AD pathology.

## 2. Methods

### 2.1 Animals

All animal experiments followed ethical and experimental procedures in agreement with Directive 2010/63 of the European Parliament and of the Council, and all the experiments in this study were preapproved by the Champalimaud Animal Welfare Body and the national competent authority (Direcção Geral de Alimentação e Veterinária, DGAV). The mouse strain used for this research project, B6;129-Tg(APPSwe,tauP301L)1Lfa Psen^1tm1Mpm^/Mmjax, RRID:MMRRC_034830-JAX, was obtained from the Mutant Mouse Resource and Research Center (MMRRC) at The Jackson Laboratory, an NIH-funded strain repository, and was donated to the MMRRC by Frank Laferla, Ph.D., University of California, Irvine.

3Tg-AD mice exhibit pathological and behavioural changes specific to AD, including deposition of extracellular amyloid beta (Aβ) plaques, intracellular neurofibrillary tangles and memory impairment, and are one of the most commonly employed mouse models for AD research [23,24]. Female 3Tg-AD and normal control (NC) mice (B6129SF2/J) were evaluated at 8, 14, 20 and 32 weeks of age to reflect the pre-, sub-, early- and mid-clinical stage of AD, respectively, as performed in a recent ASL study [25]. The evaluation consisted of behavioural testing followed by MRI. In total 40 3Tg and 31 NC mice were included in the study (N_TG,8w_ = 10, N_NC,8w_ = 10; N_TG,14w_ = 10, N_NC,14w_ = 7; N_TG,20w_ = 10, N_NC,20w_ = 7; N_TG,32w_ = 10, N_NC,32w_ = 7). All mice had free access to food and water and were housed in cages in an environmentally controlled room under a 12-h light/dark cycle.

### 2.2 Behaviour testing

Memory impairment has been observed in rodent models of Alzheimer’s disease (AD) through various behaviour tests, which can be used to assess the severity of AD symptoms [26]. Specifically, we employed the Y maze spontaneous alternations (SA) test to assess cognitive deficits and short-term memory. The maze has three identical arms which the mice are free to explore. When the animals visit three distinct arms consecutively, the pattern is referred to as a spontaneous alternation. As rodents typically prefer to investigate new environments, a decreased SA score (i.e. the percentage of SA from the total number of arm entries) might reflect a possible impairment in short term memory. Such results have been shown in transgenic 3Tg-AD mice[27].

The Y-maze was built in-house from white matte acrylic with the following dimensions (arm length: 35 cm, width: 5 cm, and height: 15 cm; angle between arms: 120 degrees) [28]. The set-up was placed in a foamed box to improve isolation from external sources of noise and vibration and was equipped with a 30 fpm video camera to record the movement and infrared lights to reduce any stressful elements from the environment [29]. The animals were handled for 20 min a day for 2 days before the behaviour experiments [26]. On the day of the testing, the mice were acclimatised to the experimental room for at least 30 minutes, after which they were introduced to the maze and left free to explore for a total of 8 minutes (2 minutes for habituation and 6 minutes for testing).

#### 2.2.1 Behavioural Data Analysis

Based on the videos, the sequence of arm entries was manually recorded. An arm entry was considered only if all four limbs of the animal were within the arm and an alternation was counted in the event of consecutive entries into three distinct arms. The SA score (%SA) was then calculated as the percentage of spontaneous alternations from the total number of arm entries:

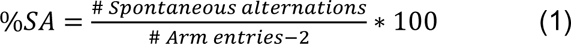

### 2.3 In-vivo MRI for mapping tissue perfusion

In this study we employed non-invasive arterial spin labelling (ASL) MRI to estimate cerebral blood flow (CBF) in three different brain regions - cortex, hippocampus (HC), and midbrain (MB). We map various quantitative parameters, including lateral ventricular volume, BCSFB-mediated total water delivery, tissue CBF and T1 (cortex, hippocampus and midbrain), as well as T1 values of CSF (T1_csf_).

Images were acquired on a 9.4 T Bruker BioSpec scanner operating an AVANCE III HD console and using a gradient system capable of producing up to 660 mT/m in each direction. An 86 mm volume was used for transmission and ensured a relatively uniform B1 profile, while a 4-element array receive-only cryogenic coil (Bruker BioSpin, Fallanden, Switzerland) was used for signal reception [30].

#### 2.3.1 Anatomical reference scans

A T2-TurboRARE sequence (fast-spin echo, Paravision v6.0.1) was applied to acquire sagittal and coronal anatomical reference images to clearly visualise the location of the major CSF compartments in the brains of control and 3Tg mice. T2-TurboRARE sequence parameters were: field of view (FOV) = 20 mm × 16 mm; matrix size = 200 × 160; RARE factor = 8; effective echo time (TE) = 35.61 ms; repetition time (TR) = 2200 ms [20].

Sagittal anatomical reference images (15 × 0.4 mm slices) were used to position the coronal anatomical reference imaging slice and the ASL imaging slices. Coronal anatomical reference images (6 × 0.4 mm slices, 2.4 mm total) were manually positioned to align with the caudal region of the lateral ventricles. Subsequently, these coronal images were manually segmented to provide subject-wise estimates of lateral ventricular volume [20].

#### 2.3.2 Standard ASL Acquisition

A flow-sensitive alternating inversion recovery (FAIR) labelling scheme was used for the acquisition of standard-ASL and BCSFB-ASL data [19,31]. The FAIR technique alternates between a global inversion (non-selective, M_c_) and a slice-selective inversion [32]. The difference between the non-selective (labelled) and slice-selective (control) images results in a perfusion-weighted ASL image that reflects the signal from the labelled blood water that has moved into the imaging slice in the given inflow time (TI). Measuring this difference at multiple inflow times enables the quantification of perfusion related parameters, such as CBF. As described in previous work, FAIR-ASL data were acquired using a single slice, single shot spin echo – echo planar imaging readout, 20 mm slice-selective width, and a global labelling pulse (non-selective)[31].

*Standard-ASL acquisition parameters:* 1 mm imaging slice thickness, matrix size = 40 × 56, FOV = 20 mm × 20 mm, 7 dummy scans, TE = 20 ms. Inflow times (TI) = [200, 500, 1000, 1500, 2000, 3000, 4000 ms], using a repetition time (TR) = 10000 ms, 5 repetitions per TI [31].

#### 2.3.3 BCSFB ASL Acquisition

By using an ultra-long echo time (TE) parameter in BCSFB-ASL, the acquisition is fine-tuned to detect the delivery of labelled arterial water across the BCSFB, into the ventricular CSF [19], due to the much longer T2 of CSF compared to parenchymal tissue. This BCSFB-ASL signal represents the total delivery of labelled water, and therefore does not provide a measure of CSF secretion, i.e. the net movement across the BCSFB. The BCSFB-ASL also employs a FAIR encoding with the same slice selective and global labelling pulses as for the standard ASL protocol.

*BCSFB-ASL acquisition parameters:* single slice, 2.4 mm imaging slice thickness, matrix size = 32 × 32, FOV = 20 mm × 20 mm, 5 dummy scans, TE = 220 ms. TI = [200, 200, 750, 1500, 2750, 4000, 5000, 6000 ms], using a recovery time (TRec) = 12000 ms, 10 repetitions per TI [21,31]. Using the previously acquired anatomical reference images, the BCSFB-ASL imaging slice was manually positioned to align with the caudal end of the lateral ventricles, as it has been shown to be densely populated with CP tissue [33]. The evaluation of BCSFB function in this study was centred on CP tissue in the lateral ventricles, excluding the 3^rd^ and 4^th^ ventricles.

### 2.4 ASL-MRI analysis

Standard-ASL and BCSFB-ASL data were analysed to obtain absolute values of tissue CBF (cortical, HC, MB) and total BCSFB-mediated water delivery, respectively, as described previously [19] (see Figure 1). For the quantification of CBF from standard-ASL data, three regions of interest (ROI) were drawn for each subject across the cerebral cortex, the hippocampus, and midbrain using the perfusion-weighted, ΔM FAIR-ASL image (see Figure 1f), and the mean voxel signal was calculated across each ROI. For each ASL image pair, the non-selective mean ROI value (M_c_) was subtracted from the slice-selective (labelled) mean ROI value to provide the perfusion-weighted signal, ΔM. Repeated measures of ΔM and M_c_ at each inflow time were averaged to provide [TI, ΔM] and [TI, M_c_] datasets.

**Figure 1.**
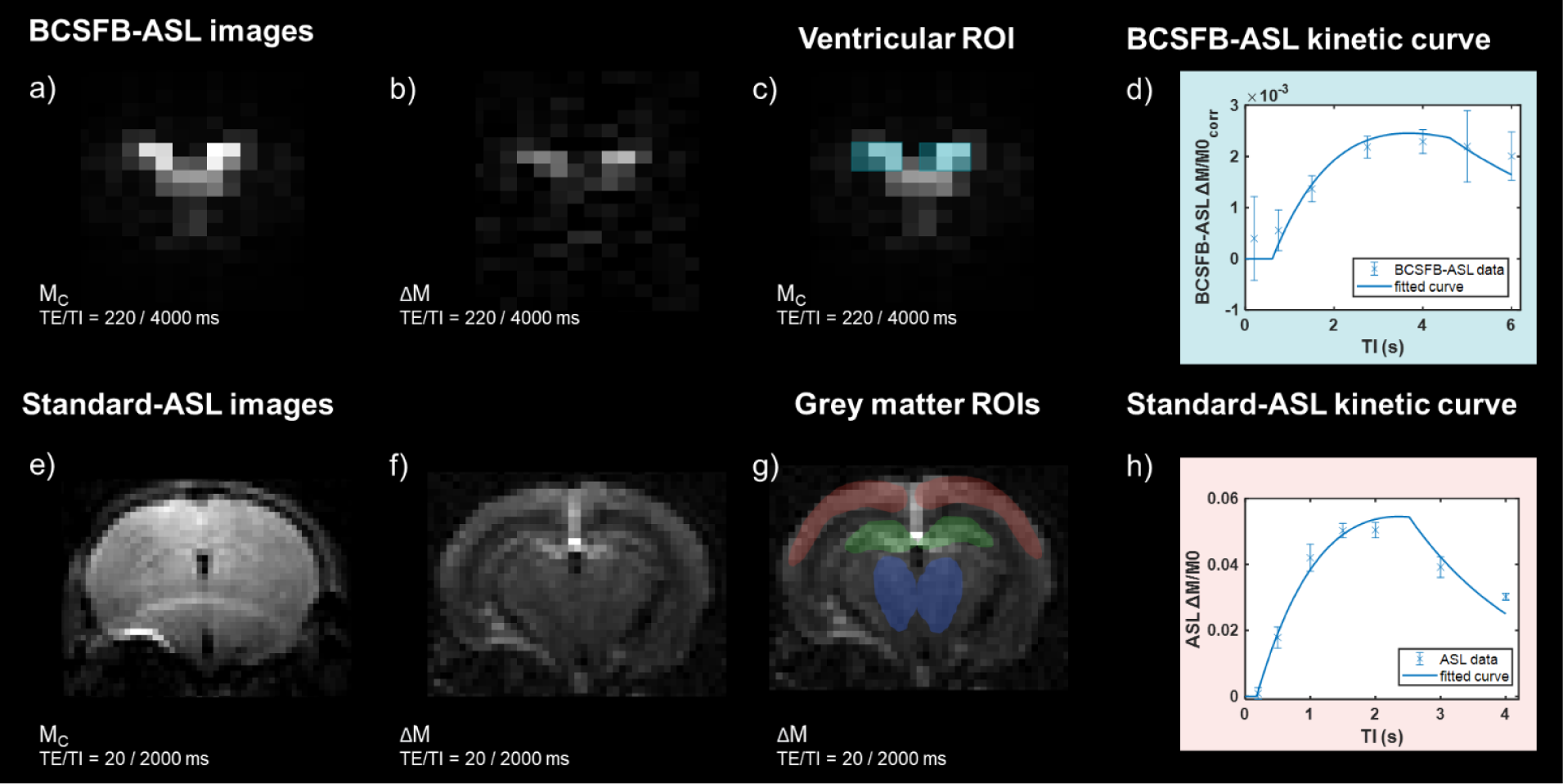
FAIR-ASL control (M_c_) and subtracted (ΔM) images for a-b) BCSBF-ASL (TE = 220 ms, TI = 4000 ms) and d-e) standard-ASL (TE = 20 ms, TI = 2000 ms). ROI selection for ASL analysis. c) BCSFB-ASL ROI selection (turquoise) for BCSFB-mediated water delivery quantification. f) standard-ASL ROI selection for CBF quantification in 3 regions: cortex (red), hippocampus (green), midbrain (blue). ASL data and fitted kinetic curves for BCSFB-ASL (d) and standard-ASL using a cortical ROI (h). Representative data from a 20w normal control mouse (N = 1, error bars: ± stdev computed over repetitions).

The [TI, M_c_] data were used to estimate the subject-wise values of M0, T1_tissue_ (M_c_ data at TE = 20 ms) and T1_csf_ (M_c_ data at TE = 220 ms) by fitting the simple inversion recovery (IR) curve described in equation 2:

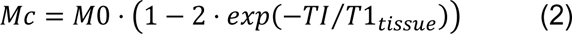

The [TI, ΔM] data were then used to fit the relevant Buxton kinetic model: a single-compartment general Buxton kinetic model for CBF from standard-ASL data [34], and a 2-compartment adaptation of the Buxton model for extracting rates of BCSFB-mediated water delivery from BCSFB-ASL data [19,31,35,36], as described in Supplementary Information. For the calculation of per-subject hemodynamic outputs, subject-wise T1 (T1_tissue_ for standard ASL or T1_CSF_ for BCSFB-ASL) and M0 (or M0_corr_ for BCSFB-ASL) values extracted from the IR fittings were used in the Buxton model as inputs when calculating CBF and BCSFB-mediated water delivery values for each subject [19].

To determine an overall measure of total lateral ventricular BCSFB-function, a summary measure of the BCSFB-ASL signal was obtained across the lateral ventricles, which permitted the measurement of the total amount of labelled arterial-blood-water delivery to ventricular CSF. Thus, for the BCSFB-ASL images, two 3 × 2 voxel ROIs (12 voxels in total, ROI volume = 11.25 mm^3^) were positioned on a non-selective (control) image, overlaid with the position of the lateral ventricles (Figure 1c). The combined ROI average signals were subtracted in a pairwise fashion to provide ΔM values. The calculated M0 will be highly dependent on ventricle size due to partial volume effects in the low resolution ASL images. A subject-wise ventricular volume normalisation factor was applied to M0_CSF_ to provide a volume-normalised, equilibrium magnetization, M0_corr_ (equation 3), which accounts for the difference between the total ventricular volume (quantified from T2-weighted anatomical images) and the ventricular ROI volume (i.e. from the 12 voxels used for the ROI from the low-resolution functional data with a volume of 11.25 mm^3^). This normalisation step is necessary for the accurate quantification of the total amount of BCSFB-mediated water delivery [19].

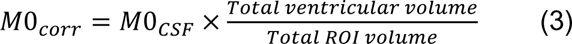

The outputs of the model fittings provided subject-wise quantitative values for CBF and T1_tissue_ (T1_cortex_, T1_HC_, T1_MB_ for standard-ASL images), and rates of BCSFB-mediated water delivery alongside T1_csf_ (BCSFB-ASL images). We report the group average of the individually extracted values.

### 2.5 Histology

Following MRI, histopathological analysis of three animals per group was used to confirm specific AD pathology. Mice were anaesthetised and transcardially perfused with phosphate buffer saline, pH 7.0 and with 4% paraformaldehyde (PFA) in accordance with protocols approved by the Champalimaud Animal Welfare & Ethical Review Body (ORBEA). Brains were collected and post-fixed for 24 h in 4% PFA. To maintain consistency of dissection and to facilitate analysis in our cohort of mice, brains were coronally sliced in register with the Allen Mouse Brain Atlas (See http://mouse.brain-map.org). Coronal slabs were paraffin embedded and serially sectioned at 4 μm. Bregma levels 0.145, −2.055, and −2.88 were selected for immunohistochemistry for Tau and Aβ [37]. Tissue sections were immunostained for Tau and Aβ using Novolink^TM^ Polymer (Leica Biosystems, UK), detected with DAB and counterstained with Harris Hematoxylin. For Tau, primary antibody (anti-Phospho-Tau, clone AT8, Thermo Fisher, cat. MN1020) was used at 1:100 dilution overnight at 4°c, after antigen retrieval with citrate buffer pH6 for 20 minutes at 98° c. For Aβ, primary antibody (anti-Beta-Amyloid, clone 6E10, Biolegend, cat. 803007) was used at 1:750 dilution overnight at 4°c, after antigen retrieval with 80% formic acid for 10 min at room temperature. Brightfield images were acquired with a Zeiss AxioScan.Z1 fully automated slide scanner using the 20x objective lenses. Finally, sections were analysed for density and spatial distribution of AT8 positive neurons and plaque distribution.

### 2.6 Statistical methods

GraphPad Prism 10 was used to conduct statistical testing. Statistical comparisons between control and 3Tg data, at each time point, were conducted using unpaired, 2-tailed, Mann-Whitney tests, after Shapiro Wilkes normality testing revealed that most parameters were not normally distributed. Given that Shapiro Wilkes normality testing revealed that the behavioural data from the Y-maze (spontaneous alternations and total entries) was normally distributed, Welch’s t-tests were employed. Importantly, for the MR imaging data, and then separately for the behavioural data, the corrected false discovery rate (FDR) method (Benjamini and Yekutieli) method was used to account for multiple comparisons, with a desired FDR of 5%.

When comparing subjects scanned at both 14-week and 42-week (Supplementary material), paired, 2-tailed t-tests were conducted to investigate longitudinal changes of the same subjects (GraphPad Prism 10), after Shapiro Wilkes normality testing revealed that most parameters were normally distributed.

A two-way ANOVA was performed to investigate the effects of ageing, genetic background (control vs AD), and their interaction, on BCSFB function. This was then followed by Sidak’s multiple comparisons testing.

We report the group mean of the data at a given time point, alongside the standard error of the mean (SEM). Overlaid onto the figures, asterisks denote the level of statistical significance achieved in our analyses: (*) represents significance at the p ≤ 0.05 level, (**) at p ≤ 0.01, (***) at p ≤ 0.001, and (****) at p ≤ 0.0001.

## 3. Results

### 3.1 ASL results

#### 3.1.1 Standard ASL

Standard-ASL data was acquired for the quantification of CBF in three gray matter regions: the cortex, hippocampus (HC) and midbrain (MB) (Figure 2, Figure S1). For the cortical ROI, Figure 2a presents the group-averaged kinetic curves, calculated by averaging ΔM/M0 across animals, which reveal a potential trend towards an increase in cortical CBF within the 3Tg group relative to the control group at all four timepoints. This increase in cortical CBF is most evident at 32 weeks. However, the comparison of individually extracted CBF values in all three ROIs (cortex, HC and MB) revealed no significant difference between the two groups at any time point, after adjusting for multiple comparisons.

- Figure 2b shows the cortical CBF data: 8w:[(control 185 ± 11, 3Tg 208 ± 5 ml/min/100g), p = 0.92], 14w:[(control: 181 ± 6, 3Tg: 206 ± 5 ml/min/100g), p = 0.17], 20w:[(control: 197 ± 9, 3Tg: 217 ± 6 ml/min/100g), p = 0.60], 32w:[(control: 167 ± 17, 3Tg: 235 ± 11 ml/min/100g), p = 0.17].
- Figure 2c shows the HC CBF data: 8w:[(control: 246 ± 6, 3Tg: 255 ± 6 ml/min/100g), p > 0.99], 14w:[(control: 235 ± 3, 3Tg: 259 ± 6 ml/min/100g), p = 0.17], 20w:[(control: 252 ± 5, 3Tg: 267 ± 8 ml/min/100g), p > 0.99], 32w:[(control: 222 ± 17, 3Tg: 257 ± 10 ml/min/100g), p > 0.99].
- Figure 2d shows the MB CBF data: 8w:[(control 258 ± 6, 3Tg 263 ± 4 ml/min/100g), p > 0.99], 14w:[(control: 231 ± 6, 3Tg: 247 ± 6 ml/min/100g), p = 0.86], 20w:[(control: 234 ± 7, 3Tg: 251 ± 4 ml/min/100g), p = 0.55], 32w:[(control: 205 ± 16, 3Tg: 222 ± 5 ml/min/100g), p > 0.99].

**Figure 2.**
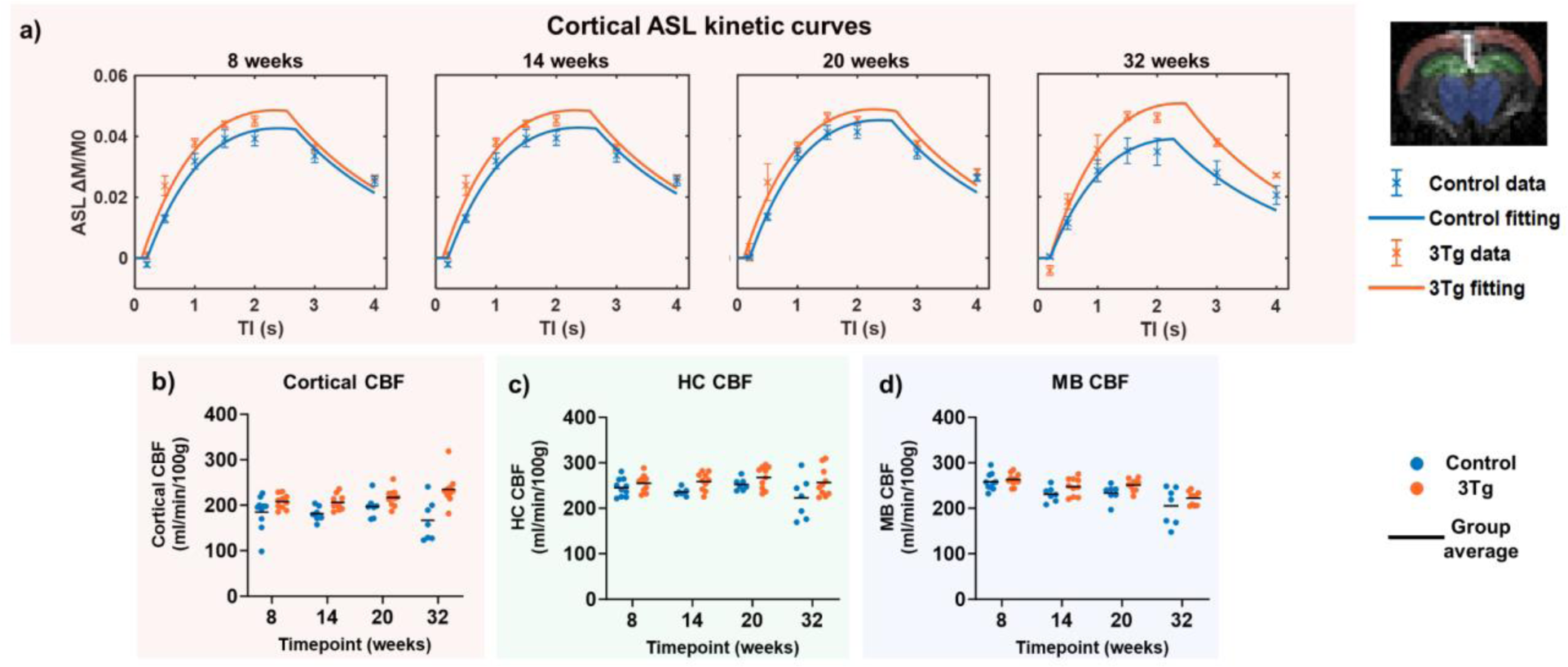
Gray matter perfusion: standard-ASL MRI data describing blood flow: a) group-averaged cortical ASL kinetic curves for control and 3Tg cohorts, and b)-d) individual subject and group averaged CBF values, at each time point for ROIs covering the cortex, hippocampus and midbrain.

T1 measurements in these regions also indicated that any differences between control and 3Tg groups were not statistically significant at any of the time points, after correcting for multiple comparisons (Figure S1).

#### 3.1.1 BCSFB ASL

BCSFB-ASL MRI data was collected to quantify total BCSBF-mediated water delivery (Figure 3). Figure 3a presents the group-averaged kinetic curves, calculated by averaging ΔM/M0_corr_ across animals, which reveal increased water delivery in the 3Tg cohort, relative to strain-matched controls, at all four timepoints, an effect which can be readily observed from visual inspection of the kinetic curves (Figure 3a). Furthermore, Figure 3b highlights the statistically significant difference in the values of the BCSFB-mediated water delivery between the two groups when comparing individual subject data: 8w:[(control 0.55 ± 0.09, 3Tg 1.13 ± 0.10 µl/min), p = 0.036], 14w:[(control: 0.80 ± 0.07, 3Tg: 1.34 ± 0.04 µl/min), p = 0.015], 20w:[(control: 0.81 ± 0.06, 3Tg: 1.39 ± 0.09 µl/min), p = 0.036], 32w:[(control: 0.74 ± 0.11, 3Tg: 1.32 ± 0.09 µl/min), p = 0.041].

**Figure 3.**
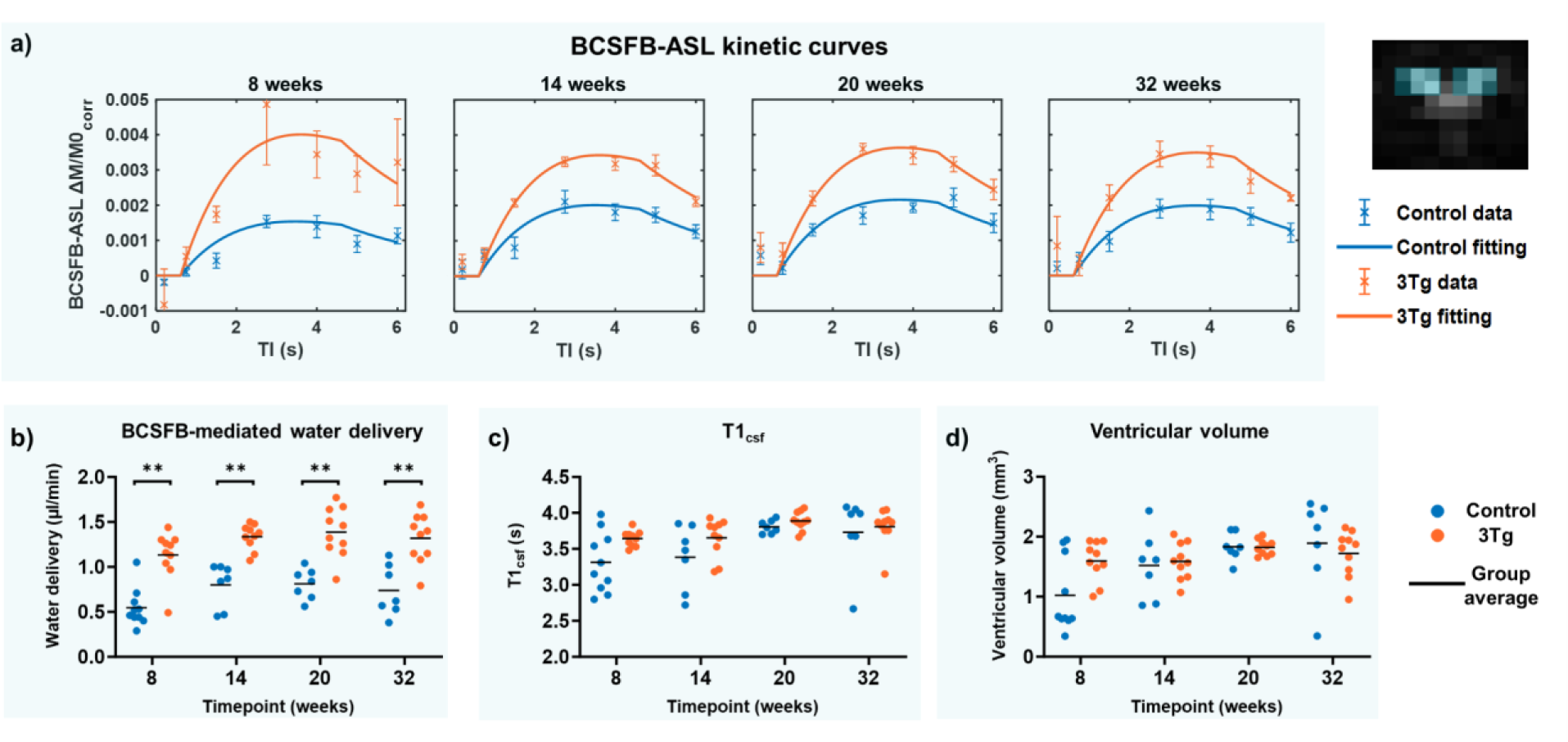
BCSFB function and ventricular homeostasis: BCSFB-ASL MRI data describing water delivery: a) group-averaged kinetic curves for control and 3Tg cohorts, and b) individual subject and group average total water delivery values, at each time point. Individual subject and group average c) T1_csf_ values from control image (Mc) data, and d) ventricular volume measurements from structural MR data.

T1_csf_ was also calculated for each subject, and was not found to be significantly different between the groups, at any time point (Figure 3c): 8w:[(control 3.31 ± 0.13, 3Tg: 3.64 ± 0.03 s), p = 0.60], 14w:[(control: 3.38 ± 0.17, 3Tg: 3.65 ± 0.08 s), p > 0.99], 20w:[(control: 3.81 ± 0.04, 3Tg: 3.89 ± 0.04 s), p > 0.99], 32w:[(control: 3.73 ± 0.19, 3Tg: 3.81 ± 0.08 s), p > 0.99].

Individual subject ventricular volume measurements revealed no significant differences between the groups, at any time points (Figure 3d): 8w:[(control 1.02 ± 0.19, 3Tg: 1.60 ± 0.11 mm3), p = 0.55], 14w:[(control: 1.52 ± 0.21, 3Tg: 1.59 ± 0.10 mm3), p > 0.99], 20w: [(control: 1.83 ± 0.09, 3Tg: 1.82 ± 0.04 mm3), p > 0.99], 32w:[(control: 1.89 ± 0.29, 3Tg: 1.72 ± 0.12 mm3), p > 0.99]. T1_csf_ at 8 and 14 weeks, and ventricular volumes at 8 weeks, displayed a trend towards an increase in the 3Tg mice, but was not found to be statistically significant after correcting for multiple comparisons.

A Two-Way ANOVA further confirmed that differences in BCSFB-mediated water delivery between control and AD subjects were significantly driven by the group effect, i.e. control or AD status (F(1, 63) = [99.07], p<0.0001). Furthermore, the effect of age/time point was shown to be a significant driver for differences in BCSFB function (F = 4.534, p = 0.0061). However, the interaction term was shown to be not significant (F= 0.04245, p = 0.99) (Table S1). Sidak’s test for multiple comparisons found that differences in the mean water delivery rate was significantly different exclusively in instances where controls were compared to 3Tg subjects, and not within the groups (Table S2).

Several 3Tg subjects scanned at 14 weeks were also rescanned at 42 weeks (N = 5), which allowed for post-hoc investigations into the effects of ageing on BCSFB function (Figure S2) and CBF (Figure S3) within the 3Tg group. No significant differences in BCSFB-mediated water delivery were observed between the two ages (Figure S2 a and b), despite there being a trend towards an increase with ageing (p = 0.10). T1_csf_ and ventricular volume measurements indicated a significant increase with ageing (Figure S2 c and d). Furthermore, HC CBF was found to be significantly decreased with ageing, with no changes in HC T1 (Figure S3 d-f). However, for cortical and MB regions, CBF and T1 values were not found to be significantly different (Figure S3 a-c, g-i).

### 3.2 Behaviour

The behaviour assessment using the spontaneous alternations task showed significant difference between the control and the 3Tg mice starting with 20 weeks of age, as illustrated in Figure 4. In particular, the number of arm entries is significantly reduced for the 3Tg animals (20w:[(control: 41 ± 8, 3Tg: 22 ± 7), p = 0.00014], 32w:[(control: 28 ± 4, 3Tg: 16 ± 6), p = 0.00019], Figure 5c). The SA score is also lower in the 3Tg mice compared to controls after 20 weeks (20w:[(control: 61 ± 8, 3Tg: 45 ± 14), p = 0.019], 32w:[(control: 63 ± 6, 3Tg: 53 ± 8), p = 0.013]), however, the differences are not statistically significant when correcting for multiple comparisons. Generally, the AD mice spent more time close to the middle of the maze, especially the older animals. We also observed a positive correlation between the SA score and the number of arm entries (r = 0.39, p = 0.01) for the 3Tg mice, while no such correlation was observed for the control animals. These behaviour results which probe the willingness of the mice to explore the maze, as well as their working memory, substantiate the known traits of the 3Tg mice which start to exhibit cognitive decline between 3- to 5- months of age [38]. Moreover, for the AD mice, a negative trend between the BCSFB water delivery and SA score is observed (r = -0.25, p = 0.11), although the correlation is not significant.

**Figure 4.**
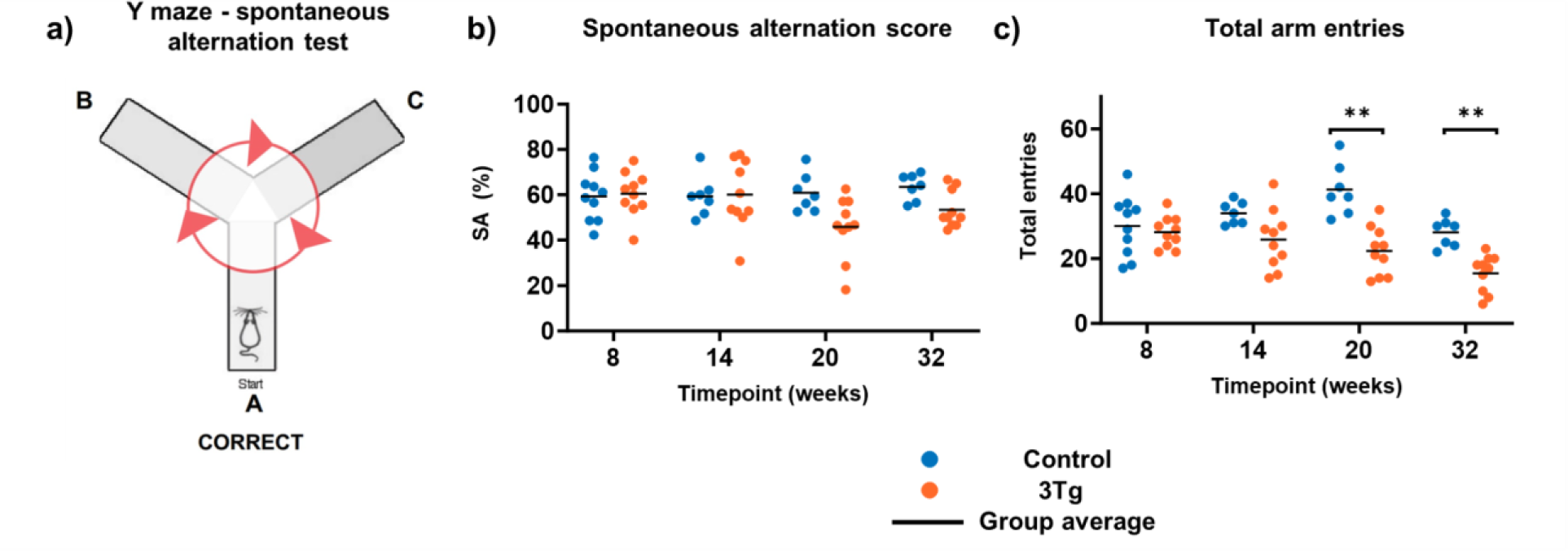
Behaviour analysis. A) Schematic representation of the Y-maze and a correct spontaneous alternation; b) spontaneous alternation score for control and 3Tg animals at different ages; c) total number of arm entries for control and 3Tg animals at different ages.

**Figure 5.**
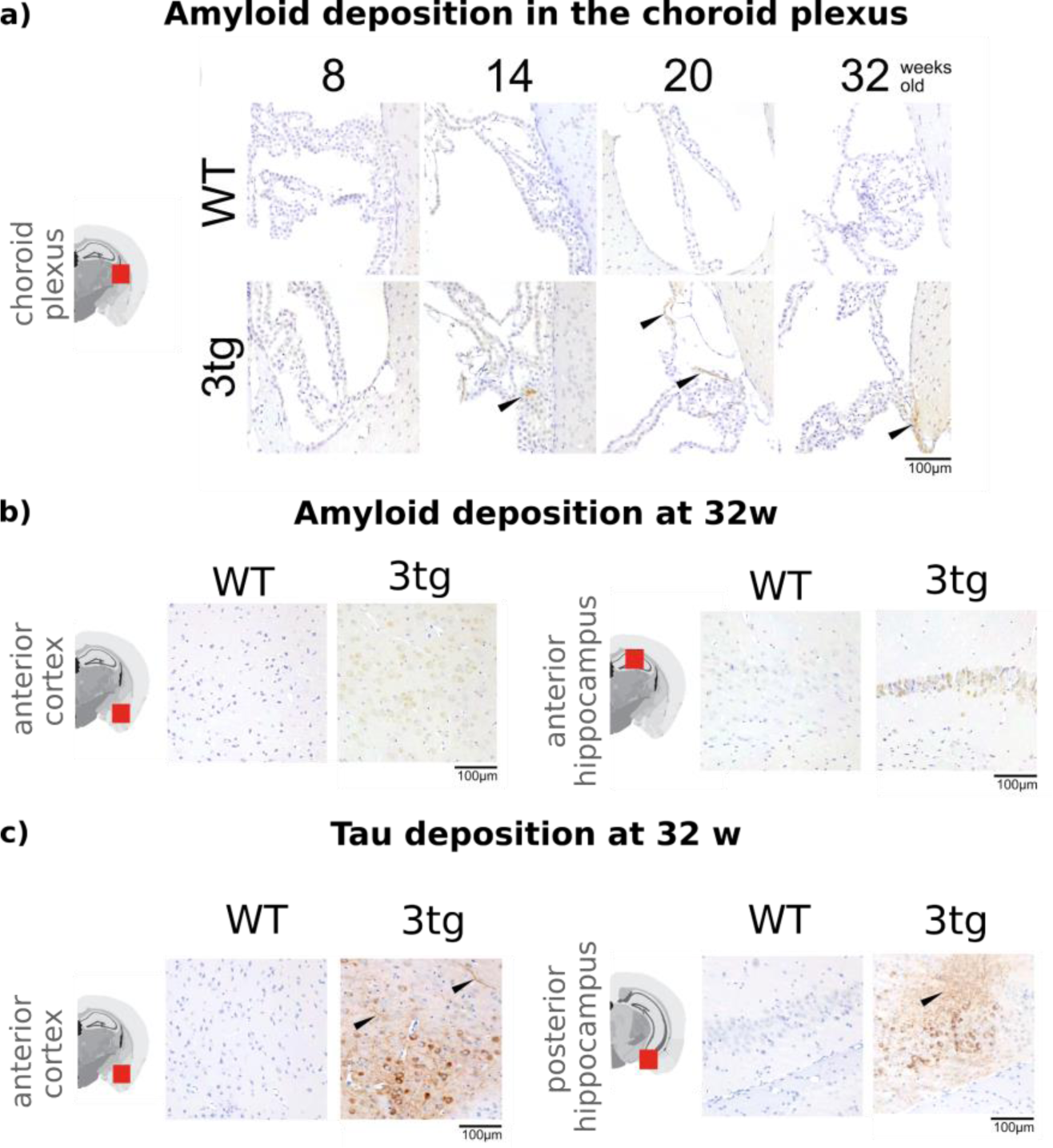
Immunohistochemical staining of Aβ and tau proteins in different areas of the brain for controls and 3Tg mice. A) Aβ staining in the choroid plexus with plaques detected starting with 14 weeks. B) Aβ staining in the hippocampus and amygdala at 32 weeks, showing wide-spread plaque formation. C) Tau staining in the hippocampus and amygdala at 32 weeks, showing intracellular deposition and neurofibrillary tau tangles (black arrowhead).

### 3.3 Histology

Immunohistochemical staining of Aβ and tau proteins confirms the expected pathology of the 3Tg mouse model of AD (Figure 5), with amyloid plaques detected in the choroid plexus starting at 14 weeks of age (Figure 5a). We observe positive staining in regions of the brain involved early in the progression of AD, such as the hippocampus and amygdala, already at 8 weeks (Figure S5), with an increase in the staining intensity with age for both proteins. We also detect neurofibrillary tau-tangles starting at 20 weeks in the hippocampus and 32 weeks in the amygdala (Figure S6). As expected, no staining is observed in the control animals.

## 4. Discussion

Our work harnessed a recently developed BCSFB-ASL MRI technique and showed, in-vivo, that CP function is altered in the early stages of Alzheimer’s disease in the widely employed triple transgenic mouse model. The measured differences in BCSFB function precede measured changes in behaviour or widespread deposition of neurofibrillary tangles, and are not accompanied by significant changes in ventricular volume or brain tissue perfusion, suggesting that, at least in this mouse model, the CP may play an important role in the early pathophysiology of AD. Given the non-invasive and translational nature of ASL MRI, measurements of BCSFB function can thus become a new and sensitive biomarker for AD.

### 4.1. BCSFB-mediated water delivery

In this work we find that the 3Tg mouse model of AD consistently exhibits an elevated degree of BCSFB-mediated labelled blood-water delivery to the ventricular CSF (Figure 3) relative to age-matched control mice. The ultra-long TE ASL measurement employed here reflects the exchange of the labelled blood water across the BCSFB, depending on the total volume of the CP in the LV convolved with its perfusion and the extraction fraction, i.e. the permeability of the BCSFB to labelled blood water. In order to further understand the observed increased water delivery in 3Tg mice, we performed ultra-high resolution ex-vivo MRI on perfused whole brain samples to measure the volume of the CP within the LV of 3Tg and control animals at 12-14 months of age. We found no significant difference in the estimated CPs volume between the groups, with mean values of 0.0782 ± 0.0089 µl and 0.0849 ± 0.0363 µl for control and 3Tg animals respectively, see Supplementary Information S5. This finding suggests that a putative difference in CP volume does not underpin the observed increase in BCSFB-mediated water delivery in this animal model and that this is therefore reflective of a functional change at the choroid plexus.

We speculate that the increased rate of blood-to-CSF water delivery likely has two components: 1) a breakdown in the BCSFB integrity, giving rise to increased water permeability in the 3Tg model, due to the many morphological changes within CP tissue [4,7], including the deposition of amyloid beta plaques (Figure 5); 2) an increase in CP perfusion for boosting clearance due to the presence of adherent proteins, an effect recently reported in AD patients, but not for MCI, although with smaller effect size compared to our results. Comparing the MRI biomarkers with the behavioural analysis did not reveal any statistically significant correlations. However, a negative trend between water delivery across BCSFB and the SA score has been observed and is an avenue for future exploration.

### 4.2 Current results in the context of known CP morphology and functional changes in AD

Investigations into CP tissue during AD in rodents have revealed that there is an impairment in function and metabolism of the BCSFB, with studies demonstrating that both the rate of secretion and the secretory profile of CSF are anomalous in the human and rodent brain during ageing and Alzheimer’s disease [4,7,39–43]. In this work, histological analysis revealed the presence of amyloid in the CP tissue of 3Tg mice, supporting previous studies reporting on the role of CP in the clearance of amyloid beta from the CNS [3,43]. For instance, previous histological analysis of mice injected with oligomeric Aβ revealed several proteins constituting cytoplasmic aggregates, including tau and amyloid. These aggregates not only mechanically disrupt plasma membranes of choroid plexus epithelial cells (Cpecs), but also facilitate a marked decline in the integrity of the BCSFB in AD through the downstream activation of matrix metalloproteases, and a downregulation of enzymatic activity and tight junction protein expression at the BCSFB locus [44]. These functional changes within the Cpec working unit of the BCSFB will be closely intertwined with changes in Cpec morphometry. Metabolic alterations [40], alongside oxidative stress – an early event in AD pathogenesis [45] – will lead to substantial Cpec death. Together, these factors will likely have a drastic effect on the transport of water across the BCSFB, through changes primarily in permeability of the BCSFB through e.g. paracellular means [44]. However, given the limited literature describing the effects of AD on the CP tissue and associated vasculature, as well as the fact that several functional parameters contribute to the BCSFB-ASL measurements, we cannot exclude the possibility of other factors increasing the measured rate of water delivery across the BCSFB in 3Tg mice, such as increased activity of transporters involved in water movement, as well as increased perfusion to the BCSFB itself.

### 4.3 Tissue perfusion

A recent meta-analysis of neuroimaging data [46] found ASL MRI measures of cerebral perfusion to show the greatest degree of ‘biomarker abnormality’ at the early stages of late onset Alzheimer’s Disease. Thus, in the present study we hypothesised that ASL measures of brain tissue perfusion (here taken in the cortex, hippocampus and midbrain) would differ between the 3Tg model and controls. However, although a trend of increased CBF in the 3Tg was observed at selected timepoints (see Figure 2), accounting for multiple comparisons, we found no significant difference between the 3Tg and control group in our ASL measures of tissue perfusion within the ROIs examined across the timepoints. To our knowledge this is the first report of cerebral perfusion measurements in the 3Tg model. This observation is broadly consistent with previous findings [47] where no difference in whole brain uptake of 18F-FDG was found between 11-month 3Tg and control mice. However, other studies report hypometabolism of 18F-FDG in the 16 month 3Tg [48] which may be expected to be accompanied by a concomitant reduction in perfusion as assessed with ASL-MRI.

### 4.4 Ageing effects on water delivery across BCSFB

Post-hoc analyses were conducted to determine the extent to which subject age was playing a role in the differences observed in our hemodynamic biomarkers. From the Two-Way ANOVA, both the effect of genetic background and ageing were shown to be significant drivers, with no significant interaction term. Following multiple comparison corrections, significant differences in BCSFB-mediated water delivery were observed only when comparing control with 3Tg groups at any time point, and not intra-group differences at different timepoints. Furthermore, paired comparisons within the 3Tg cohort, from 14 to 42 weeks, indicated a trend towards increased rates of BCSFB-mediated water delivery. The lack of statistical significance here may be a result of the small sample size for this analysis (N = 5). However, when taken together, our findings suggest that the underlying AD pathology is indeed the primary driver for the observed changes in BCSFB water delivery, regardless of the specific time point.

### 4.5 Limitations and future work

There are several limitations to this study. For example, this work clearly demonstrates a derangement in BCSFB function in the 3Tg mouse model of AD compared to age-matched controls. However, it is important to note that the BCSFB-ASL measurements reflect the convolution of the total CP volume with its perfusion and the extraction fraction. To disentangle these effects, future studies could also include measurements of CP perfusion, which are highly challenging due to the required resolution. Nevertheless, several steps have been taken in this direction both in mice [49] and in humans [50] by acquiring ASL data with high spatial resolution and/or multiple echo times. We can further leverage an innovative bed design for acquiring high resolution ASL data in rodents that we recently proposed [51], as well as denoising strategies for the analysis [52].

MRI measurements were non-invasively collected in the anaesthetised mouse brain, but not the awake brain. Anaesthetic protocols are known to variably affect cerebral hemodynamics in rodent strains, which can limit the accuracy of quantification of hemodynamic parameters such as CBF [53,54] relative to the ground truth under normal awake physiology. However, awake imaging is highly challenging in mice, with further concerns regarding the impact of stress on hemodynamic markers [55,56]. Thus, the use of a standardised protocol which prioritises high scan-rescan reproducibility [19] fast reversibility and recovery [54,57], as well as straightforward administration, was favourable in this work.

Another limiting factor is the fact that the main study design was cross-sectional, thus we are unable to evaluate subject-wise longitudinal changes in the hemodynamic biomarkers, except for a small animal cohort, which would provide further aetiological understanding of disease onset, and should be considered in future work.

The scope of the histological analysis in this study was mainly focused on confirming the presence of known pathology. The limited sample size (N = 3 per group) and spatial coverage are insufficient for quantitative comparisons between the in-vivo biomarkers and brain-wide burden of plaques and tangles. Future work could benefit from experiments designed to understand the underlying tissue and mechanistic changes that lead to differences in water delivery across BCSFB.

### 4.6 Summary and clinical perspectives

Contributing to the recent efforts in understanding the critical role of the CP-BCSFB locus in AD pathology, this study shows a significant increase in BCSFB-mediated water delivery in the 3Tg mouse model of AD, for the first time. Our findings are also in line with recent clinical studies showing an increase in apparent CP perfusion in AD [14,58]. The far greater effect size of the BCSFB-ASL measurements relative to traditional ASL measurements suggests that the BCSFB-ASL method may prove to be a promising readily translatable biomarker of early AD pathology.

## Acknowledgements

The authors acknowledge the vivarium of the Champalimaud Center for the Unknown, a facility of CONGENTO financed by Lisboa Regional Operational Programme (Lisboa 2020), project LISBOA01-0145-FEDER-022170. CP and JW are supported by the Wellcome Trust (225345/Z/22/Z). AI and RC are supported by “la Caixa” Foundation (ID 100010434) and from the European Union’s Horizon 2020 research and innovation programme under the Marie Skłodowska-Curie grant agreement No 847648, fellowship code CF/BQ/PI20/11760029. DLT is supported by the UCLH NIHR Biomedical Research Centre.

The authors have no relationships/activities/interests related to the content of the manuscript to disclose.

## Supporting Information

### S1 – ASL modelling

For standard ASL, cerebral blood flow (CBF) for the cortex, HC, and MB, was quantified by taking the ASL signal, i.e. ΔM values at TE = 20 ms, and fitting the general kinetic Buxton model (single compartment model), as described by equation 3 from Buxton et al 1998 [34].

For BCSFB-ASL, rates of delivery of labelled blood water to ventricular CSF were quantified using a 2-compartment perfusion model to the ASL signal (TE = 220 ms). This 2-compartment model was first described by Alsop and Detre [35], and then later adapted by Wong *et al*. [59]. Subsequently, this model was further adapted by Evans *et al.* for the purposes of describing the transfer of labelled water from blood into the ventricular CSF, rather than into the cerebral cortex [19] (Equation 4, Figure 7).

When fitting the ASL curves we assume an inversion efficiency α of 0.9, and a known value for the longitudinal relaxation of arterial blood (T1_b_ **=** 2.5 s, from the literature) [60]. We also assume a constant blood-to-CSF water partition coefficient ϕ, which is the ratio of the density of water in the blood to the density of water in the CSF, and as such, is taken to have a value of 1 [19,49,61,62].

Curve Fitting Toolbox (Matlab) was used throughout this work to fit kinetic curves to the acquired multi-TI standard-ASL and BCSFB-ASL data to the models described, using in-house scripts [19–21,31].

### S2 – Standard ASL across brain regions

**Figure S1.**
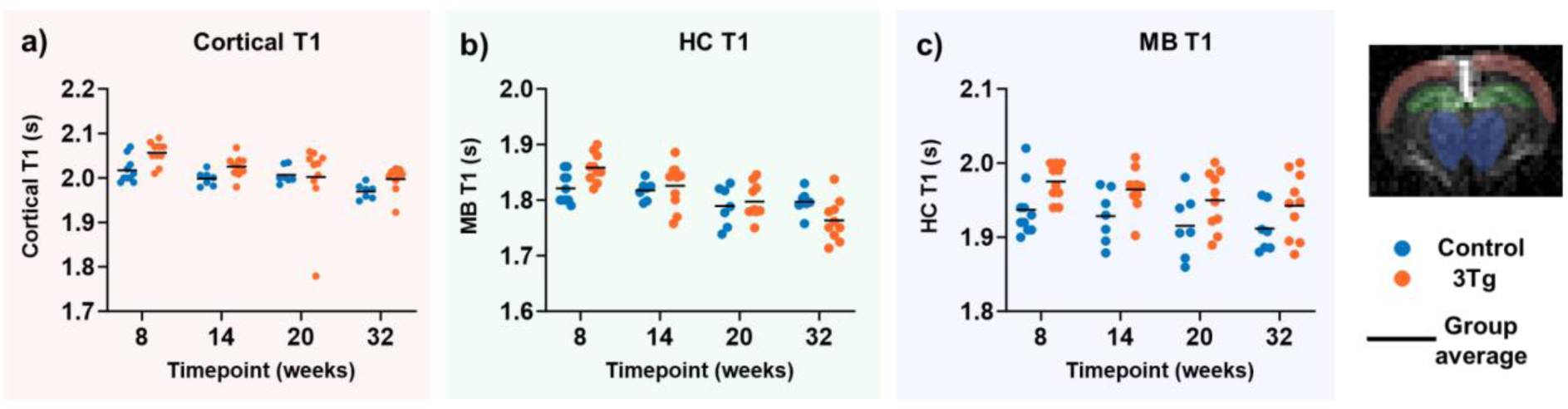
T1 in gray matter ROIs. Individual subject T1 values extracted from control image (Mc) standard-ASL data, at different time points, in a) the cortex, b) the hippocampus (HC), and the midbrain (MB).

### S3 – Effects of age, genetic background and their interaction

**Table S1:** Summary of Two-Way ANOVA: BCSFB-mediated water delivery. A Two-Way ANOVA was conducted to investigate the effects of ageing (time point), genetic background (control vs 3Tg status), and their interaction, on BCSFB-mediated water delivery rates. SS: sum of squares, DF: degrees of freedom, MS: mean squares.

| Source of variation | SS (Type III) | DF | MS | F | P value |
| --- | --- | --- | --- | --- | --- |
| Interaction | 0.007187 | 3 | 0.002396 | F = 0.04245 | P=0.9882 |
| Time point | 0.7677 | 3 | 0.2559 | F = 4.534 | P=0.0061 |
| Genetic background | 5.591 | 1 | 5.591 | F = 99.07 | P<0.0001 |
| Residual | 3.555 | 63 | 0.05643 |  |  |
| Total | 9.920887 | 70 |  |  |  |

**Table S2:** Post-hoc multiple comparisons results from Two-Way ANOVA: BCSBF-mediated water delivery. Following the initial Two-Way ANOVA to investigate the effects of ageing and genetic background on BCSBF function, post-hoc, Sidak’s multiple comparisons testing was conducted.

| Šídák's multiple comparisons test | Summary | Adjusted P Value |
| --- | --- | --- |
| 8w control vs. 8w 3Tg | **** | <0.0001 |
| 8w control vs. 14w control | ns | 0.6108 |
| 8w control vs. 14w 3Tg | **** | <0.0001 |
| 8w control vs. 20w control | ns | 0.5254 |
| 8w control vs. 20w 3Tg | **** | <0.0001 |
| 8w control vs. 32w control | ns | 0.9562 |
| 8w control vs. 32w 3Tg | **** | <0.0001 |
| 8w 3Tg vs. 14w control | ns | 0.1549 |
| 8w 3Tg vs. 14w 3Tg | ns | 0.8325 |
| 8w 3Tg vs. 20w control | ns | 0.1977 |
| 8w 3Tg vs. 20w 3Tg | ns | 0.4289 |
| 8w 3Tg vs. 32w control | * | 0.0342 |
| 8w 3Tg vs. 32w 3Tg | ns | 0.9246 |
| 14w control vs. 14w 3Tg | *** | 0.0007 |
| 14w control vs. 20w control | ns | >0.9999 |
| 14w control vs. 20w 3Tg | *** | 0.0001 |
| 14w control vs. 32w control | ns | >0.9999 |
| 14w control vs. 32w 3Tg | ** | 0.0011 |
| 14w 3Tg vs. 20w control | *** | 0.0009 |
| 14w 3Tg vs. 20w 3Tg | ns | >0.9999 |
| 14w 3Tg vs. 32w control | **** | <0.0001 |
| 14w 3Tg vs. 32w 3Tg | ns | >0.9999 |
| 20w control vs. 20w 3Tg | *** | 0.0002 |
| 20w control vs. 32w control | ns | >0.9999 |
| 20w control vs. 32w 3Tg | ** | 0.0016 |
| 20w 3Tg vs. 32w control | **** | <0.0001 |
| 20w 3Tg vs. 32w 3Tg | ns | >0.9999 |
| 32w control vs. 32w 3Tg | *** | 0.0002 |

### S4 – Longitudinal analysis in a sub-cohort of animals

A sub-cohort of N = 5 AD mice from the 14 weeks group was re-evaluated at 42 weeks, following the same procedure including behaviour testing followed by ASL MRI.

The results presented in Figure S2 show that the BCSFB water delivery kinetic curve at 42 weeks is above the one from 14 weeks, with a significant increase in ventricular volume and T1_CSF_ and a trend towards an increase in the water delivery parameter. The standard ASL results presented in Figure S3 reveal a decrease in CBF measurements in hippocampus, while the other regions and parameters do not show significant differences between ages.

**Figure S2.**
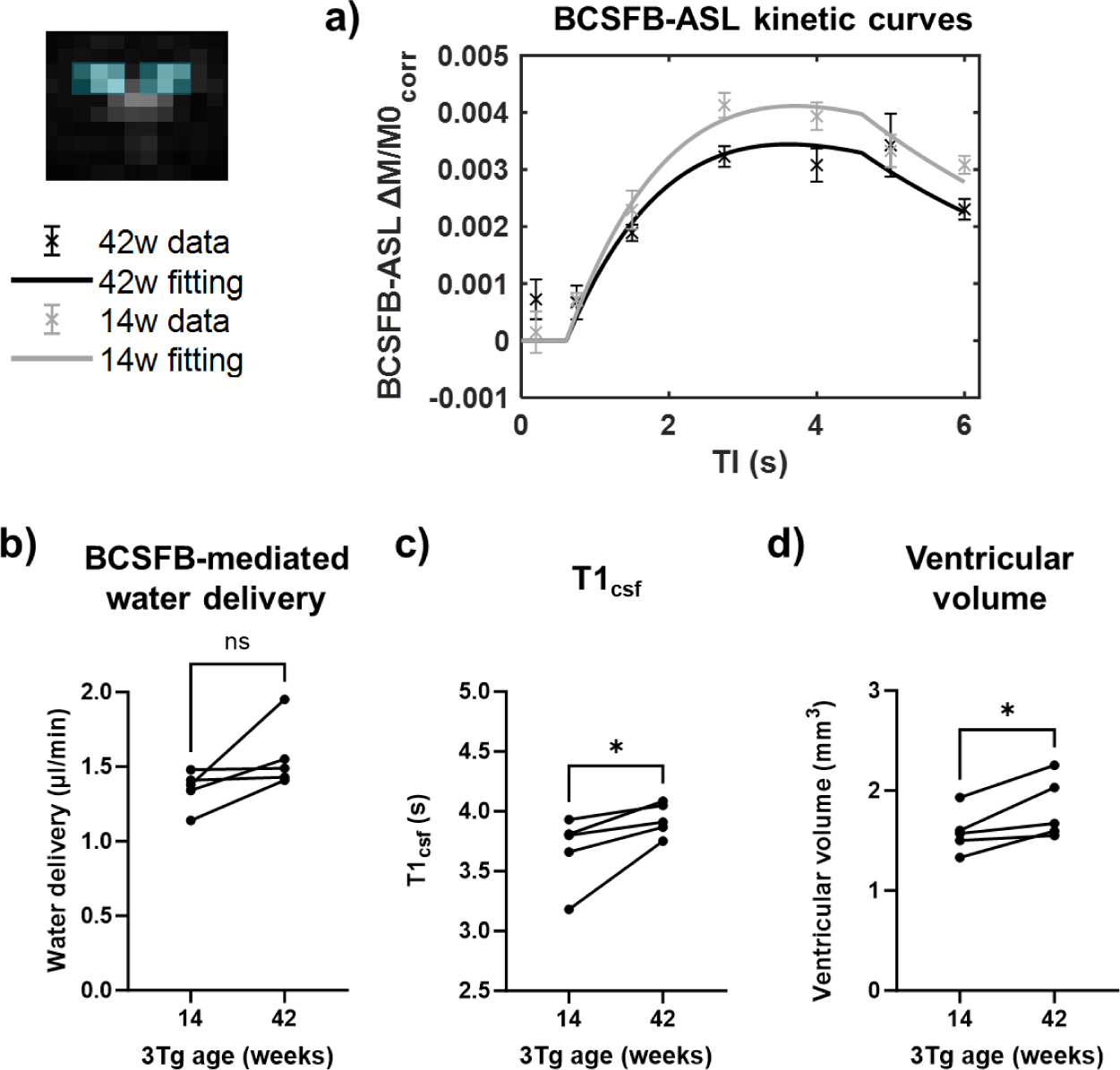
Longitudinal comparison of BCSFB function and ventricular homeostasis. BCSFB-ASL MRI data describing water delivery in 14 vs 42 weeks 3Tg subjects (longitudinal, N = 5). A) group-averaged kinetic curves, and b) individual subject and group average total water delivery values, at each time point. Individual subject c) T1csf values from control image (Mc) data, and c) ventricular volume measurements from structural MR data.

**Figure S3.**
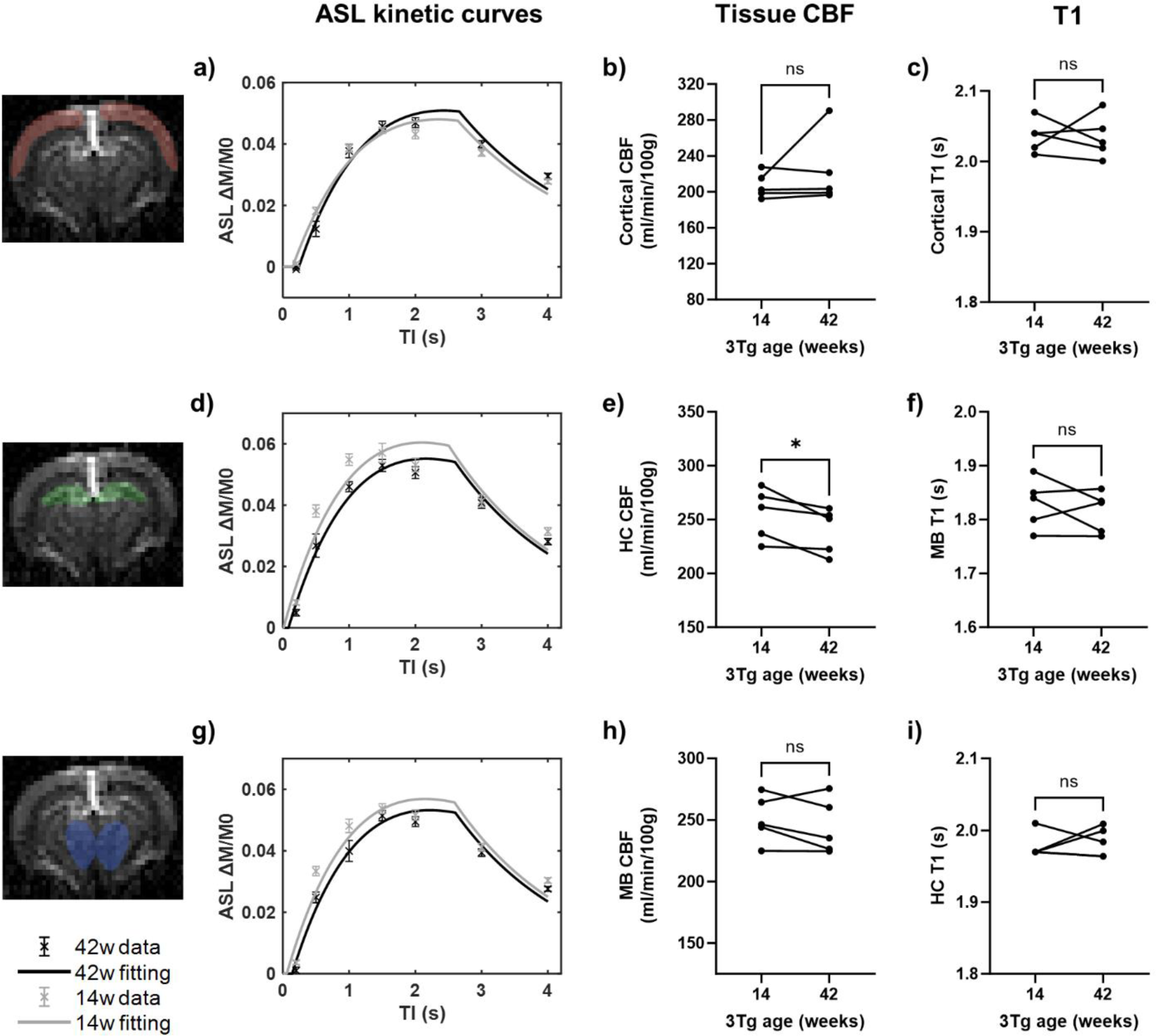
Longitudinal comparison of CBF and T1. Standard-ASL MRI data describing blood flow in 14 vs 42 weeks 3Tg subjects (longitudinal, N = 5). Top row: cortical region of interest (ROI), middle row: hippocampus (HC) ROI, bottom row: midbrain (MB) ROI. Group-averages kinetic curves for 14 vs 42 weeks (a,d,g), individual subject CBF values (b, e, h), and individual subject T1 values from control image (Mc) data (c, f, i).

### S5 – Ex-vivo microimaging of choroid plexus

To assess the volume of the choroid plexus we performed ultra-high resolution MRI of ex-vivo brain tissue, comparing control and AD mice.

#### S5.1 Sample preparation

For ex-vivo MRI, female mice (N_NC_ = 3, mean age 12.5 months and N_TG_ = 4, mean age 14 months) were anaesthetised and transcardially perfused with 4% PFA. The brains were surgically removed, post-fixed for 24 h in 4% PFA and rehydrated in PBS for at least 24h [63]. Before being mounted in 10mm NMR tubes filled with fluorinert, the brains were submerged in fluorinert and the cerebral hemispheres were slightly separated with a flexible spatula, so that fluorinert enters the ventricular system. In this way, we ensured an excellent contrast to directly image and quantify the volume of the choroid plexus.

#### S5.2 Image acquisition

Ultra-high resolution data at 40 μm isotropic was acquired on a 16.4T Aeon Ascent Bruker Scanner equipped with a Micro 5 imaging probe capable of generating 3T/m gradients. Images were acquired using a 3D gradient echo (GE) sequence with the following parameters: TE/TR = 5/45 ms, matrix size = 450 x 225 x 255, flip angle = 12°, NA = 12 averages, with a total scan duration of ∼8h per sample. MR microscopy images used to quantify the choroid plexus are shown for a representative sample in Figure S4 a-c.

#### S5.3 Data analysis

The volume of the choroid plexus was calculated from the histogram analysis of the ventricular signal. Following the manual delineation of the lateral ventricles, including the background signal and choroid plexuses, the histogram of the signals was analysed to quantify the volume of the choroid plexus while accounting for partial volume in the imaging voxels. First, the main peak of the distribution, that corresponds to the background signal (noise) in areas filled with fluorinert, was fitted with a Rayleigh distribution describing the MRI noise in the absence of signal, as illustrated in Figure S4d. Next, to determine the tissue signal, the fitted Rayleigh distribution was subtracted from the original histogram. The resulting histogram was rescaled between a lower signal boundary (LB) corresponding to a voxel with no tissue and an upper signal boundary (UB) corresponding to a voxel with 100% tissue. We calculated LB as the mean of the Rayleigh distribution, and UB as two noise standard deviations below the maximum measured signal. Then, to account for partial volume effects, the CP volume was calculated from the histogram count weighted by the signal value.

The CP volume estimated for different animals is shown in Figure S4, with a mean value of 0.0782 ± 0.0089 µl for the control group and 0.0849 ± 0.0363 µl for the AD group. We do not observe significant differences between the normal controls and AD mice.

**Figure S4.**
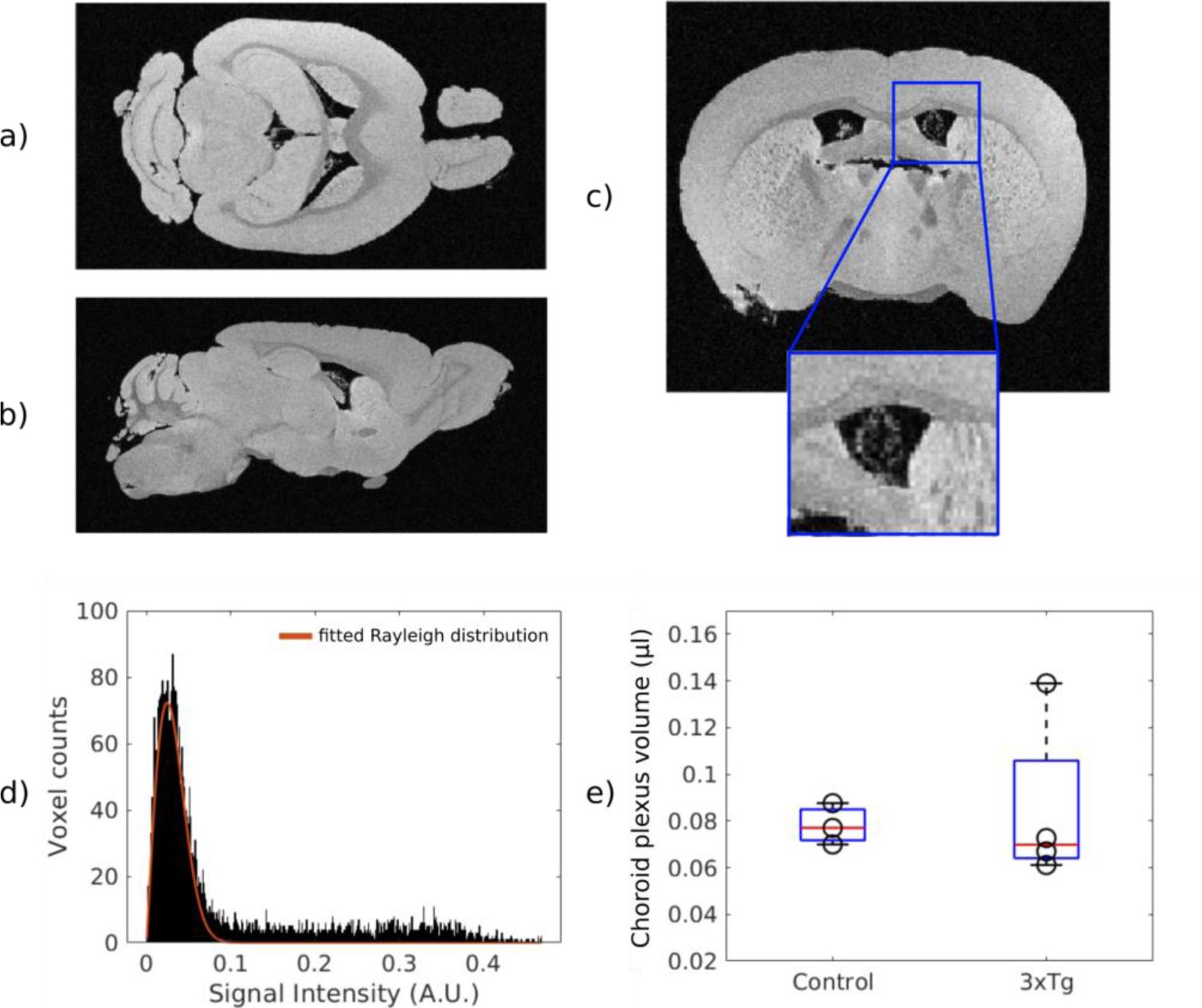
a-c) Example of MR microscopy images for one control mouse in the horizontal, sagittal and coronal planes, with an inset showing the clear contrast of the choroid plexus which appears as bright voxels inside the ventricles. D) Distribution of signal intensity values inside the ventricles and fitted Rayleigh distribution. E) Boxplots showing the median and interquartile range of CP volume for the controls and 3Tg animals.

### S6 – Histology: Aβ and tau staining at different ages

**Figure S5.**
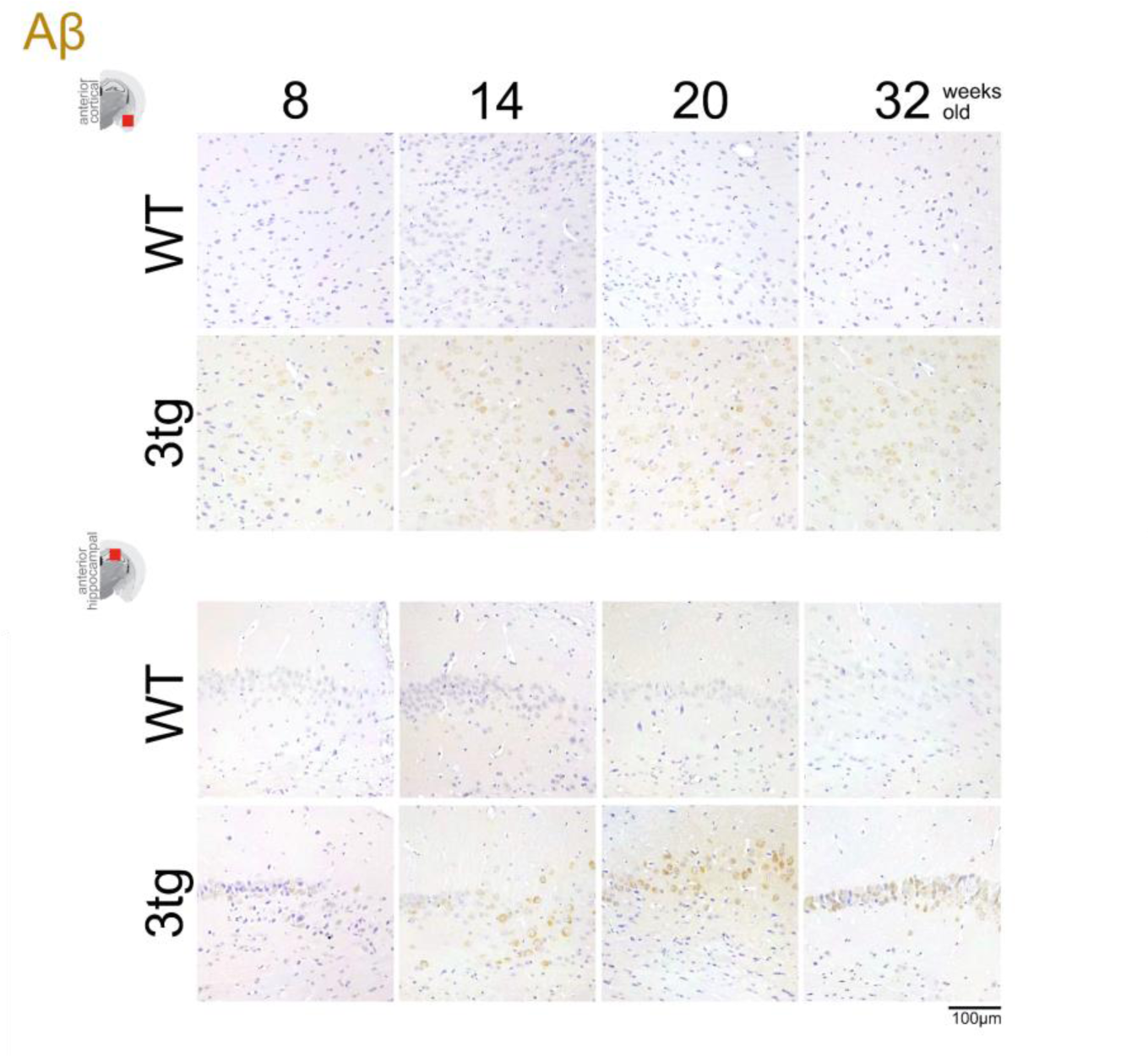
Immunohistochemical staining of amyloid beta in the anterior cortical (top) and hippocampal (bottom) regions for wild type and 3Tg mice of different ages. In the 3Tg animals, plaques are detected already ay 8 weeks, although the staining becomes more pronounced after 14 weeks. No plaques are detected in the control animals.

**Figure S6.**
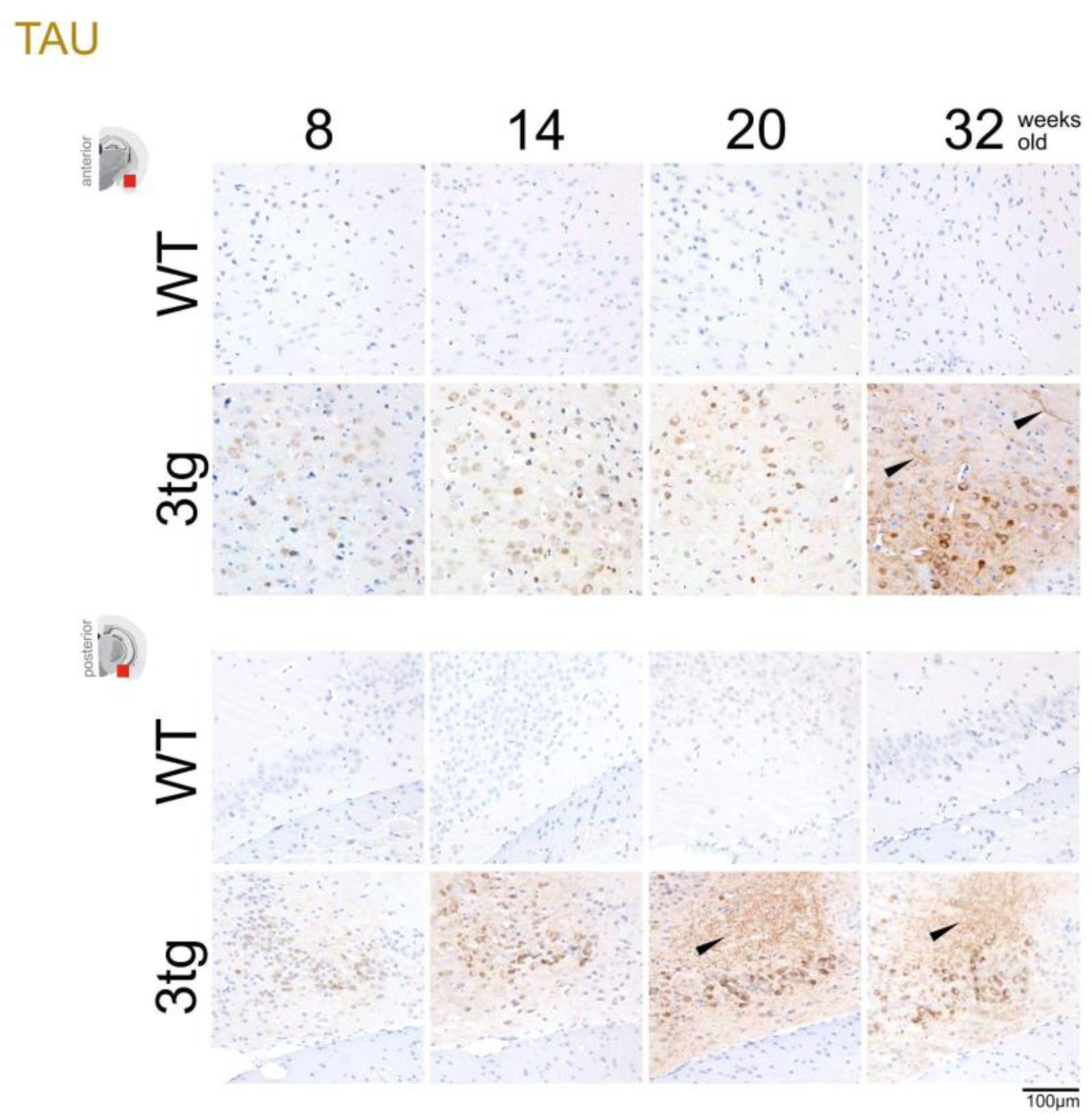
Immunohistochemical staining of tau in the anterior cortical (top) and posterior hippocampal (bottom) regions for wild type and 3Tg mice of different ages. In the 3Tg animals, positive tau stains are detected already ay 8 weeks, and neurofibrillary tau tangles (arrowheads) are observed at 20 weeks and 32 weeks in the hippocampus and cortical regions, respectively. No tau staining is detected in the control animals.

## References

1. Faraci FM, Mayhan WG, Farrell WJ, Heistad DD. Humoral regulation of blood flow to choroid plexus: role of arginine vasopressin. Circ Res. 1988;63: 373–379.

2. Alvira-Botero X, Carro EM. Clearance of amyloid-β peptide across the choroid plexus in Alzheimer’s disease. Curr Aging Sci. 2010;3: 219–229.

3. Elbert DL, Patterson BW, Lucey BP, Benzinger TLS, Bateman RJ. Importance of CSF-based Aβ clearance with age in humans increases with declining efficacy of blood-brain barrier/proteolytic pathways. Commun Biol. 2022;5: 98.

4. Balusu S, Brkic M, Libert C, Vandenbroucke RE. The choroid plexus-cerebrospinal fluid interface in Alzheimer’s disease: more than just a barrier. Neural Regeneration Res. 2016;11: 534–537.

5. Peng W, Achariyar TM, Li B, Liao Y, Mestre H, Hitomi E, et al. Suppression of glymphatic fluid transport in a mouse model of Alzheimer’s disease. Neurobiol Dis. 2016;93: 215–225.

6. Nedergaard M, Goldman SA. Glymphatic failure as a final common pathway to dementia. Science. 2020;370: 50–56.

7. Marques F, Sousa JC, Sousa N, Palha JA. Blood-brain-barriers in aging and in Alzheimer’s disease. Mol Neurodegener. 2013;8: 38.

8. Bouwman FH, Frisoni GB, Johnson SC, Chen X, Engelborghs S, Ikeuchi T, et al. Clinical application of CSF biomarkers for Alzheimer’s disease: From rationale to ratios. Alzheimers Dement. 2022;14: e12314.

9. Serot JM, Béné MC, Foliguet B, Faure GC. Morphological alterations of the choroid plexus in late-onset Alzheimer’s disease. Acta Neuropathol. 2000;99: 105–108.

10. Stopa EG, Tanis KQ, Miller MC, Nikonova EV, Podtelezhnikov AA, Finney EM, et al. Comparative transcriptomics of choroid plexus in Alzheimer’s disease, frontotemporal dementia and Huntington’s disease: implications for CSF homeostasis. Fluids Barriers CNS. 2018;15: 18.

11. González-Marrero I, Giménez-Llort L, Johanson CE, Carmona-Calero EM, Castañeyra-Ruiz L, Brito-Armas JM, et al. Choroid plexus dysfunction impairs beta-amyloid clearance in a triple transgenic mouse model of Alzheimer’s disease. Front Cell Neurosci. 2015;9: 17.

12. Choi JD, Moon Y, Kim H-J, Yim Y, Lee S, Moon W-J. Choroid Plexus Volume and Permeability at Brain MRI within the Alzheimer Disease Clinical Spectrum. Radiology. 2022;304: 635–645.

13. Delvenne A, Tijms BM, Gobom J, Redolfi A, Barkhof F, Zetterberg H, et al. Choroid plexus volume is associated with levels of CSF proteins predominantly expressed by the choroid plexus in non-demented individuals with AD pathophysiology. Alzheimers Dement. 2022;18. doi:10.1002/alz.067517

14. Lu P, Li J, Zhao L. Increased apparent blood flow and volume of the choroid plexus in the Alzheimer’s Disease patients. Alzheimers Dement. 2023;19. doi:10.1002/alz.066276

15. Li Y, Rusinek H, Butler T, Glodzik L, Pirraglia E, Babich J, et al. Decreased CSF clearance and increased brain amyloid in Alzheimer’s disease. Fluids Barriers CNS. 2022;19: 21.

16. Schubert JJ, Veronese M, Marchitelli L, Bodini B, Tonietto M, Stankoff B, et al. Dynamic 11C-PiB PET Shows Cerebrospinal Fluid Flow Alterations in Alzheimer Disease and Multiple Sclerosis. J Nucl Med. 2019;60: 1452–1460.

17. de Leon MJ, Li Y, Okamura N, Tsui WH, Saint-Louis LA, Glodzik L, et al. Cerebrospinal Fluid Clearance in Alzheimer Disease Measured with Dynamic PET. J Nucl Med. 2017;58: 1471–1476.

18. Muthuraman M, Oshaghi M, Fleischer V, Ciolac D, Othman A, Meuth SG, et al. Choroid plexus imaging to track neuroinflammation - a translational model for mouse and human studies. Neural Regeneration Res. 2023;18: 521–522.

19. Evans PG, Sokolska M, Alves A, Harrison IF, Ohene Y, Nahavandi P, et al. Non-Invasive MRI of Blood–Cerebrospinal Fluid Barrier Function. Nat Commun. 2020;11: 1–11.

20. Perera C, Harrison IF, Lythgoe MF, Thomas DL, Wells JA. Pharmacological MRI with Simultaneous Measurement of Cerebral Perfusion and Blood-Cerebrospinal Fluid Barrier Function using Interleaved Echo-Time Arterial Spin Labelling. Neuroimage. 2021;238: 118270.

21. Perera C, Tolomeo D, Baker RR, Ohene Y, Korsak A, Lythgoe MF, et al. Investigating changes in blood-cerebrospinal fluid barrier function in a rat model of chronic hypertension using non-invasive magnetic resonance imaging. Front Mol Neurosci. 2022;15: 964632.

22. Petitclerc L, Hirschler L, Wells JA, Thomas DL, van Walderveen MAA, van Buchem MA, et al. Ultra-long-TE arterial spin labeling reveals rapid and brain-wide blood-to-CSF water transport in humans. Neuroimage. 2021;245: 118755.

23. Jankowsky JL, Zheng H. Practical considerations for choosing a mouse model of Alzheimer’s disease. Mol Neurodegener. 2017;12: 89.

24. Onos KD, Sukoff Rizzo SJ, Howell GR, Sasner M. Toward more predictive genetic mouse models of Alzheimer’s disease. Brain Res Bull. 2016;122: 1–11.

25. Guo Y, Li X, Zhang M, Chen N, Wu S, Lei J, et al. Age- and brain region-associated alterations of cerebral blood flow in early Alzheimer’s disease assessed in AβPPSWE/PS1ΔE9 transgenic mice using arterial spin labeling. Mol Med Rep. 2019;19: 3045–3052.

26. Miedel CJ, Patton JM, Miedel AN, Miedel ES, Levenson JM. Assessment of Spontaneous Alternation, Novel Object Recognition and Limb Clasping in Transgenic Mouse Models of Amyloid-β and Tau Neuropathology. J Vis Exp. 2017. doi:10.3791/55523

27. Zhou L, Huang J-Y, Zhang D, Zhao Y-L. Cognitive improvements and reduction in amyloid plaque deposition by saikosaponin D treatment in a murine model of Alzheimer’s disease. Exp Ther Med. 2020;20: 1082–1090.

28. Prieur EAK, Jadavji NM. Assessing Spatial Working Memory Using the Spontaneous Alternation Y-maze Test in Aged Male Mice. Bio Protoc. 2019;9: e3162.

29. Kraeuter A-K, Guest PC, Sarnyai Z. The Y-Maze for Assessment of Spatial Working and Reference Memory in Mice. Methods Mol Biol. 2019;1916: 105–111.

30. Baltes C, Radzwill N, Bosshard S, Marek D, Rudin M. Micro MRI of the mouse brain using a novel 400 MHz cryogenic quadrature RF probe. NMR Biomed. 2009;22: 834–842.

31. Perera P. Development and Application of MRI Techniques for Non-invasive Assessment of Blood-cerebrospinal Fluid Barrier Function. UCL (University College London); 2023.

32. Kim SG. Quantification of relative cerebral blood flow change by flow-sensitive alternating inversion recovery (FAIR) technique: application to functional mapping. Magn Reson Med. 1995;34: 293–301.

33. Lein ES, Hawrylycz MJ, Ao N, Ayres M, Bensinger A, Bernard A, et al. Genome-wide atlas of gene expression in the adult mouse brain. Nature. 2007;445: 168–176.

34. Buxton RB, Frank LR, Wong EC, Siewert B, Warach S, Edelman RR. A general kinetic model for quantitative perfusion imaging with arterial spin labeling. Magn Reson Med. 1998;40: 383–396.

35. Alsop DC, Detre JA. Reduced transit-time sensitivity in noninvasive magnetic resonance imaging of human cerebral blood flow. J Cereb Blood Flow Metab. 1996;16: 1236–1249.

36. Wang J, Alsop DC, Li L, Listerud J, Gonzalez-At JB, Schnall MD, et al. Comparison of quantitative perfusion imaging using arterial spin labeling at 1.5 and 4.0 Tesla. Magn Reson Med. 2002;48: 242–254.

37. Oh K-J, Perez SE, Lagalwar S, Vana L, Binder L, Mufson EJ. Staging of Alzheimer’s pathology in triple transgenic mice: a light and electron microscopic analysis. Int J Alzheimers Dis. 2010;2010. doi:10.4061/2010/780102

38. Roda AR, Esquerda-Canals G, Martí-Clúa J, Villegas S. Cognitive Impairment in the 3xTg-AD Mouse Model of Alzheimer’s Disease is Affected by Aβ-ImmunoTherapy and Cognitive Stimulation. Pharmaceutics. 2020;12. doi:10.3390/pharmaceutics12100944

39. Redzic ZB, Preston JE, Duncan JA, Chodobski A, Szmydynger-Chodobska J. The Choroid Plexus-Cerebrospinal Fluid System: From Development to Aging. Current Topics in Developmental Biology. Academic Press; 2005. pp. 1–52.

40. Gião T, Teixeira T, Almeida MR, Cardoso I. Choroid Plexus in Alzheimer’s Disease-The Current State of Knowledge. Biomedicines. 2022;10. doi:10.3390/biomedicines10020224

41. Kant S, Stopa EG, Johanson CE, Baird A, Silverberg GD. Choroid plexus genes for CSF production and brain homeostasis are altered in Alzheimer’s disease. Fluids Barriers CNS. 2018;15: 34.

42. Riisøen H. Reduced prealbumin (transthyretin) in CSF of severely demented patients with Alzheimer’s disease. Acta Neurol Scand. 1988;78: 455–459.

43. Silverberg GD, Heit G, Huhn S, Jaffe RA, Chang SD, Bronte-Stewart H, et al. The cerebrospinal fluid production rate is reduced in dementia of the Alzheimer’s type. Neurology. 2001;57: 1763–1766.

44. Brkic M, Balusu S, Van Wonterghem E, Gorlé N, Benilova I, Kremer A, et al. Amyloid β Oligomers Disrupt Blood-CSF Barrier Integrity by Activating Matrix Metalloproteinases. J Neurosci. 2015;35: 12766–12778.

45. Nunomura A, Perry G, Aliev G, Hirai K, Takeda A, Balraj EK, et al. Oxidative damage is the earliest event in Alzheimer disease. J Neuropathol Exp Neurol. 2001;60: 759–767.

46. Dang C, Wang Y, Li Q, Lu Y. Neuroimaging modalities in the detection of Alzheimer’s disease-associated biomarkers. psychoradiology. 2023;3: kkad009.

47. Adlimoghaddam A, Snow WM, Stortz G, Perez C, Djordjevic J, Goertzen AL, et al. Regional hypometabolism in the 3xTg mouse model of Alzheimer’s disease. Neurobiol Dis. 2019;127: 264–277.

48. Stojakovic A, Chang S-Y, Nesbitt J, Pichurin NP, Ostroot MA, Aikawa T, et al. Partial Inhibition of Mitochondrial Complex I Reduces Tau Pathology and Improves Energy Homeostasis and Synaptic Function in 3xTg-AD Mice. J Alzheimers Dis. 2021;79: 335–353.

49. Lee H, Ozturk B, Stringer MS, Koundal S, MacIntosh BJ, Rothman D, et al. Choroid plexus tissue perfusion and blood to CSF barrier function in rats measured with continuous arterial spin labeling. Neuroimage. 2022;261: 119512.

50. Zhao L, Taso M, Dai W, Press DZ, Alsop DC. Non-invasive measurement of choroid plexus apparent blood flow with arterial spin labeling. Fluids Barriers CNS. 2020;17: 58.

51. Monteiro SP, Hirschler L, Barbier EL, Figueiredo P, Shemesh N. Six-fold enhancement in spatial-resolution of Pseudo-Continuous Arterial Spin Labeling Perfusion Mapping using a Cryogenic Coil at 9.4T.

52. Pires Monteiro S, Pinto J, Chappell MA, Fouto A, Baptista MV, Vilela P, et al. Brain perfusion imaging by multi-delay arterial spin labeling: Impact of modeling dispersion and interaction with denoising strategies and pathology. Magn Reson Med. 2023;90: 1889–1904.

53. Zuurbier CJ, Emons VM, Ince C. Hemodynamics of anesthetized ventilated mouse models: aspects of anesthetics, fluid support, and strain. Am J Physiol Heart Circ Physiol. 2002;282: H2099–105.

54. Munting LP, Derieppe MPP, Suidgeest E, Denis de Senneville B, Wells JA, van der Weerd L. Influence of different isoflurane anesthesia protocols on murine cerebral hemodynamics measured with pseudo-continuous arterial spin labeling. NMR Biomed. 2019;32: e4105.

55. Gao Y-R, Ma Y, Zhang Q, Winder AT, Liang Z, Antinori L, et al. Time to wake up: Studying neurovascular coupling and brain-wide circuit function in the un-anesthetized animal. Neuroimage. 2017;153: 382–398.

56. Steiner AR, Rousseau-Blass F, Schroeter A, Hartnack S, Bettschart-Wolfensberger R. Systematic Review: Anesthetic Protocols and Management as Confounders in Rodent Blood Oxygen Level Dependent Functional Magnetic Resonance Imaging (BOLD fMRI)- Part B: Effects of Anesthetic Agents, Doses and Timing. Animals (Basel). 2021;11. doi:10.3390/ani11010199

57. Hohlbaum K, Bert B, Dietze S, Palme R, Fink H, Thöne-Reineke C. Severity classification of repeated isoflurane anesthesia in C57BL/6JRj mice-Assessing the degree of distress. PLoS One. 2017;12: e0179588.

58. Alisch JSR, Kiely M, Triebswetter C, Alsameen MH, Gong Z, Khattar N, et al. Characterization of Age-Related Differences in the Human Choroid Plexus Volume, Microstructural Integrity, and Blood Perfusion Using Multiparameter Magnetic Resonance Imaging. Front Aging Neurosci. 2021;13: 734992.

59. Wong EC, Buxton RB, Frank LR. A theoretical and experimental comparison of continuous and pulsed arterial spin labeling techniques for quantitative perfusion imaging. Magn Reson Med. 1998;40: 348–355.

60. Dobre MC, Uğurbil K, Marjanska M. Determination of blood longitudinal relaxation time (T1) at high magnetic field strengths. Magn Reson Imaging. 2007;25: 733–735.

61. Chappell MA, McConnell FAK, Golay X, Günther M, Hernandez-Tamames JA, van Osch MJ, et al. Partial volume correction in arterial spin labeling perfusion MRI: A method to disentangle anatomy from physiology or an analysis step too far? Neuroimage. 2021;238: 118236.

62. Herscovitch P, Raichle ME. What is the correct value for the brain--blood partition coefficient for water? J Cereb Blood Flow Metab. 1985;5: 65–69.

63. Schilling KG, Grussu F, Ianus A, Hansen B, Barrett RLC, Aggarwal M, et al. Recommendations and guidelines from the ISMRM Diffusion Study Group for preclinical diffusion MRI: Part 2 -- Ex vivo imaging. arXiv [physics.med-ph]. 2022. Available: http://arxiv.org/abs/2209.13371

